# Comparison of CRISPR and marker based methods for the engineering of phage T7

**DOI:** 10.1101/2020.01.12.903492

**Authors:** Aurelija M. Grigonyte, Christian Harrison, Paul R. MacDonald, Ariadna Montero-Blay, Matthew Tridgett, John Duncan, Antonia P. Sagona, Chrystala Constantinidou, Alfonso Jaramillo, Andrew Millard

## Abstract

With the recent rise in interest in using lytic bacteriophages as therapeutic agents, there is an urgent requirement to understand their fundamental biology to enable the engineering of their genomes. Current methods of phage engineering rely on homologous recombination, followed by a system of selection to identify recombinant phages. For bacteriophage T7, the host genes *cmk* or *trx* have been used as a selection mechanism along with both type I and II CRISPR systems to select against wild-type phage and enrich for the desired mutant. Here we systematically compare all three systems; we show that the use of marker-based selection is the most efficient method and we use this to generate multiple T7 tail fiber mutants. Furthermore, we found the type II CRISPR-Cas system is easier to use and generally more efficient than a type I system in the engineering of phage T7. These results provide a foundation for the future, more efficient engineering of bacteriophage T7.

## 1. Introduction

Bacteriophages (phages) are viruses that specifically infect bacteria. Much of the original research on bacteriophages has guided our current understanding of molecular biology. Recently, there has been a renaissance in phage research due to the emergence of multi-drug resistant bacteria and the need for viable antibiotic alternatives[1]. Thus, there has been considerable research in the area of phage therapy, to kill bacterial infections using phages[2–4]. Phages offer a number of advantages (bactericidal agents, rapid discovery, undergo replication at the site of infection) over traditional antibiotics[5–9]. However, there are several limitations to phage therapy, such as narrow host range and limited pharmacokinetics leading to insufficient build-up of phage concentration at the site of infection[10,11].

The genetic engineering of phages offers the possibility of overcoming these limitations and allowing phage therapy to be a viable alternative. For the effective engineering of phages to occur a highly efficient and reproducible system is required. From the onset of phage research a number of methods have been used to create phage mutants; this has included random mutagenesis using a variety of inducing agents including UV, as well as chemical mutants such as hydroxylamine and N-methyl-N’-nitro-N-nitrosoguanidine[12]. The use of random mutagenesis provides the ability to create a large library of mutants, but requires extensive screening to identify a mutant of interest. Thus, other approaches have been developed to create targeted mutants using homologous recombination to introduce the desired mutation, followed by selection of the resulting phage[13]. The selection process has proven to be the major bottleneck in the creation of phage mutants. Unlike bacteria where an antibiotic resistance marker can be used to select for mutants, no such generic approach works so efficiently for phages, as phage lysis kills the bacterial host. Therefore, a number of other strategies have been developed for engineering and creating phage mutants.

An alternative to the antibiotic resistance markers used for positive selection of phage mutants, is the use of a bacterial host gene that is essential for phage replication but not essential for host growth [14,15]. In the infection of *E. coli* by phage T7, the proteins thioredoxin and cytidine monophosphate kinase (dCMP), encoded by *trxA* and *cmk* genes respectively, are known to be essential[14]. Therefore, these genes have previously been used as markers to positively select for phage mutants in an *E. coli* background that lack *trxA* or *cmk*[14,16]. This marker-based approach has been used to knockout a number of T7 genes including genes 0.4, 11, 12, 17[16–18]. These knockouts were achieved by replacing the target gene with *trxA* by homologous recombination, and selection on *E. coli* BW25113 Δ*trxA* cells [16–18].

An alternative to positive selection is to reduce the number of phage that have to be screened to find the desired mutant by increasing the efficiency of homologous recombination. An example of this is the BRED system (bacteriophage recombineering with electroporated DNA) that has been used to genetically modify mycobacteriophages and coliphages[19,20]. BRED exploits the mycobacterial recombineering system, in which expression of the RecE/RecT-like proteins of the mycobacteriophage Che9c confers high levels of homologous recombination and facilitates simple allelic exchange using a linear DNA substrate. This method has been used to engineer both lytic and temperate phages [21,22].

Another method to reduce the number of phage that require screening is the use of a Yeast Artificial Chromosome (YAC) in *Saccharomyces cerevisiae*[19,20]. Here, a phage genome can be cloned into a YAC, either in one piece or as PCR fragments assembled through gap-repair cloning [23]. Transformation-assosciated transformation (TAR) recombineering systems of yeast can then be used to introduce desired mutations, with the engineered genome re-introduced into its bacterial host. TAR approach has been used to make tail fiber mutants in coliphages T7 and T3 [24].

The most recent advance in screening methods is the use of CRISPR-Cas to counter select against phages that have not been altered, and enrich for the phage mutants of interest. A number of different CRISPR systems have now been used to modify the genomes of a number of phage systems, the first of which used a type I CRISPR system to target and create mutants of bacteriophage T7[25]. A different type I CRISPR system has also been used to create mutants of the lytic phage ICP1 that infects *V. cholera* [26]. A number of phages have been modified with type II CRISPR systems; including phage 2972 infecting *Streptococcus thermophiles*, phage P2 infecting *Lactococcus. lactis*, phiKpS2 infecting *Klebsiella pneumoniae* and phages T2, T4, T7 and KF1 that infect *E. coli* respectively [25,27–31]. A type III CRISPR system has also been used to engineer phages that infect *Staphylococcus aureus* and *Staphylococcus epidermidis* [32]. Whilst CRISPR is being used for editing phage genomes, there is still much to be learned concerning what the most efficient system to use is. For example phage T4 has been engineered by three different groups, with contrasting efficiency for type II CRISPR [29,30,33].

Both Type I and II CRISPR/Cas systems have different essential components that are required for their function. Whilst both systems require a DNA repeat-spacer, each of them has additional requirements. Type II-A CRISPR-Cas needs Cas9 endonuclease and trans-activating CRISPR RNA (*tracr*RNA), whilst the I-E CRISPR-Cas system requires Cas3 enzyme as well as a five protein complex that consists of CasA, CasB, CasC, CasD, and CasE, often referred to as cascade[34–36]. It is clear both CRISPR-Cas and marker-based selection can be used to generate phage mutants; however, no comprehensive analysis of their comparative efficiency has been carried out in a single system for a single phage.

The aim of this work was to compare efficiencies of both CRISPR-Cas systems and marker-based (*cmk* and *trx*) methods for the development of an efficient system of constructing phage T7 mutants (Figure 1). As a proof of concept, the most efficient method was then used to rapidly generate tail fiber mutants of phage T7.

**Figure 1.**
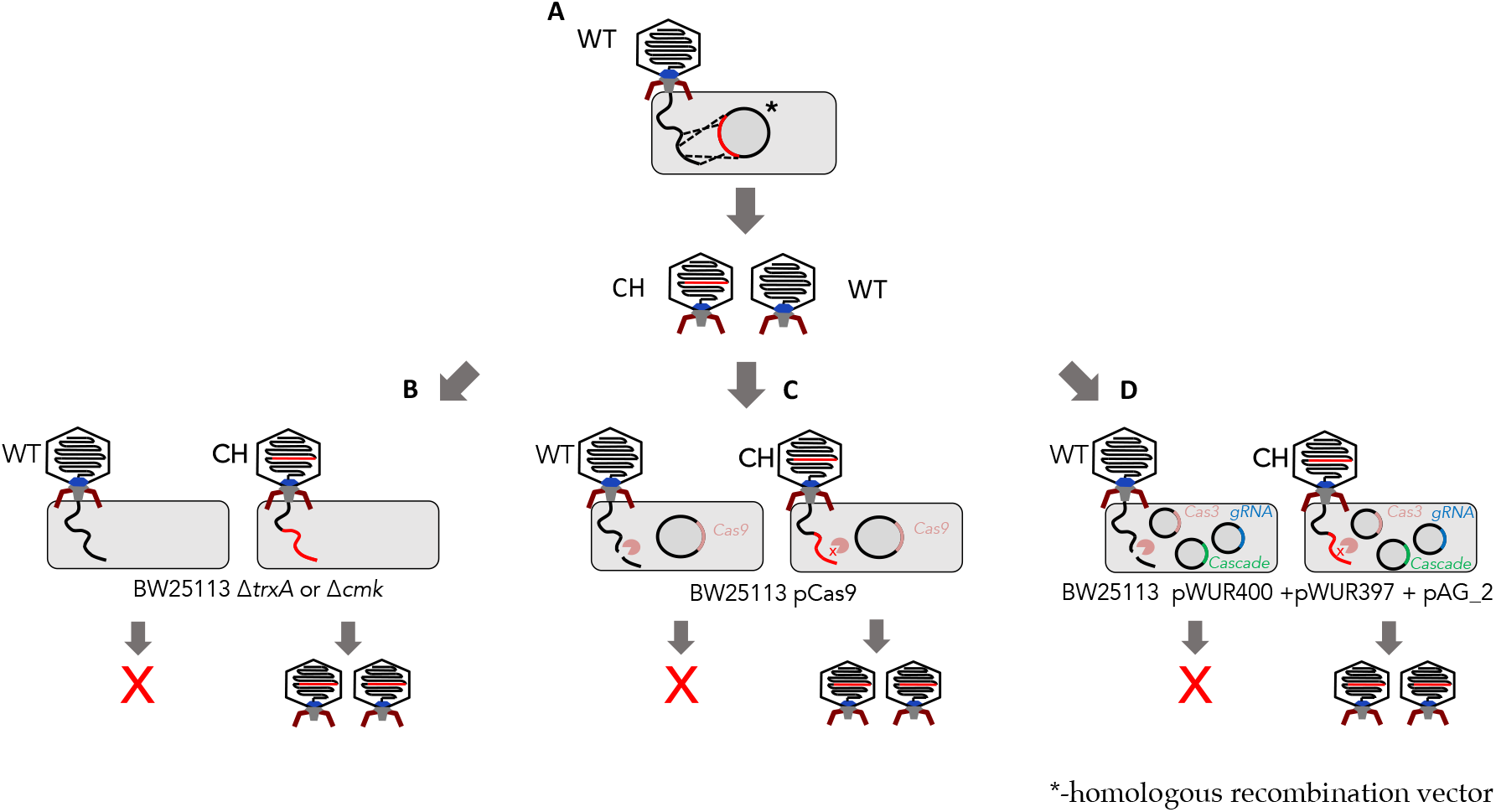
Summary of marker based vs marker-less selection methods. Homologous recombination step, allows for generation of a mixed population of chimeric/mutant (CH) and wild type (WT) phages (A). Marker based selection requires a gene encoding an essential host factor to be incorporated into the region that will homologously recombine in step A. Mutants are then selected for on *E. coli* cells that are deficient in this host factor. For phage T7 either *E.coli ΔtrxA or E.coli Δcmk* can be used for positive selection (B). Type I CRISPR system (C) consists of one vector (pCas9: contains Cas9 as well as direct repeat regions and guide RNA). A Type II CRISPR system consists of three vectors (pWUR400:Cas3, pWUR397:cascade system and pAG_2: guide RNA). Both CRISPR systems target wild-type phage, enriching for mutants.

## 2. Materials and Methods

### 2.1. Bacterial strains and Phages

*E. coli* BW25113, *E. coli* BW25113 Δ*trxA*, *E. coli* BW25113 Δ*cmk*, BL21(DE3) and BL21(AI) strains were used as the bacterial hosts for all assays carried out in this study. The strains were grown in LB medium (Oxoid, Basingstoke, UK) at 37 °C overnight with shaking (200 rpm). Phage T7 was propagated by inoculating *E. coli* culture at an OD 600 nm of 0.3. Phage lysates were filtered using a 0.22 μm pore size filter (Sartorius, Dublin, Ireland) and stored at 4 ◦C. Phage enumeration was carried out using a standard double agar overlay plaque assay or spot assays as previously described [37].

### 2.2 Type I CRISPR gRNA delivery vector component/vector design

The vectors pWUR400 and pWUR397 provide the cascade and cas proteins of the type I system [36]. A third component of the system, a vector containing a gRNA to target specific sites of interest, was designed and synthesized in a pSMART Amp^r^ vector (Fig S1). The initial vector pAG_1 contained a non-targeting gRNA control, referred to as a scrambled RNA (scr). The main components of the construct were: a) T7 RNA polymerase promoter at the 5’ of the construct; b) 5’ handle; c) gRNA for a target gene sequence; d) BbsI restriction enzyme sites flanking the gRNA region; e) 3’ handle; f) T7 terminator.

### 2.3. Type I CRISPR gRNA design

Type I CRISPR gRNAs were designed manually taking into consideration protospacer adjacent motif (PAM) requirements for the system. Previous research has identified that the PAM that fits most efficiently is AGG [38]. The main principle for the design was to first identify the AGG sequence; the 32 bp following AGG comprise the target sequence. All AGG motifs were identified in g17. Ten out of 37 identified potential gRNAs were chosen for gRNA design: four targeting the 5‘ region and six targeting the 3‘ region of the gene respectively (Figure S1). They were designed and synthesized as a complementary single stranded DNA fragments (IDT) and inserted into pAG_1 vector using Golden Gate cloning (Tables S2 and S3).

### 2.4. Type II CRISPR gRNA design

Each of the type II CRISPR gRNAs was designed using DNA 2.0 (Autumn). The following criteria were chosen when using the software: NGG was entered for the desired PAM and species off target was *E. coli*. Each of the gRNAs generated by the software were appended with BsaI enzyme restriction sites. A total of 11 gRNAs was designed and then synthesised as a complementary single stranded DNA fragments (IDT) and inserted into pCas9 vector using Golden Gate cloning (Table S2).

### 2.5. Golden Gate Cloning

The type I CRISPR gRNAs were inserted in the vector pAG1 using a modified Golden Gate cloning method [39]. Briefly, 0.5 μL of each primer (100 mM) was mixed together with 49 μL of sterile water and incubated at 95 °C for 5 minutes, prior to being diluted 10 times. The 15 μL assembly mix was prepared as follows: vector of interest (100 ng), 2 μL of the diluted primers (10 mM), 1.5 μL of 10X T4 buffer (NEB), T4 Ligase (NEB), 0.15 μL of 100x BSA and made up to 15 μL with water. The assembly mix was then incubated in a thermocycler under the following conditions: 25 cycles of 3 minutes incubation at 37 °C followed by 4 minutes at 16 °C, and 1 cycle of 5 minutes at 50 °C and 5 minutes at 80 °C. The samples were then left overnight at room temperature and 1 μL of the assembly mix was then transformed into electro-competent *E. coli* cells using 2 mm cuvettes at 2.5 kV.

### 2.6. PCR

Polymerase Chain Reaction (PCR) was carried out in 0.2 mL volume PCR in a thermal cycler (T100 Thermal cycler, Biorad, USA). HF Phusion Master mix (M0531S, New England Bio Labs, UK) and BSA (B9000S, New England Bio Labs, UK labs) were used for the reactions. Final concentrations were used as follows: primers at 0.5 μM, DNA at < 250 ng, 1x Phusion Master Mix. PCR amplification was carried out as per Protocol Phusion® High-Fidelity PCR Master Mix recommended conditions.

### 2.7 Gibson Assembly

A master mix was prepared of 5X ISO buffer 0.5 M Tris-HCl pH 7.5, 0.05 M MgCl_2_, 4 mM dNTPs (1 mM of each: dGTP, dCTP, 320 μL dATP, dTTP), 0.05 M DTT, 0.25 % (v/v) PEG800 and 5 mM NAD. Then the following components were mixed 0.26% 5X ISO Buffer, 0.005 U T5 exonuclease, 0.03 U Phusion polymerase, 5.3 U Taq ligase. The Gibson assembly was based on previously described method [40]. Briefly, fragments required for the assembly were PCR amplified (using primers in Table S3) and gel purified. Gibson reactions were carried out in 20 μL volumes, with 15 μL of Gibson master mix (0.26% 5X ISO Buffer, 0.005 U T5 exonuclease, 0.03 U Phusion polymerase, 5.3 U Taq ligase) with the remaining 5 μL made up of vector and inserts at the appropriate ratios. For the assembly of three or more fragments an equilmolar ratio of fragments was used; when inserting one fragment into a vector, a ratio of 1:2 (vector to insert) was used. The reaction was incubated at 50°C for 60 min, followed by incubation at 37 °C for 60 min. The reaction volume was diluted 3-fold and 1 μL used in electroporation.

### 2.8 Homologous recombination and in trans complementation

A strain containing a homologous recombination plasmid (HR) was grown until it reached an OD600 of 0.3-0.4, whereby phage T7 was then added at MOI ~0.01. This was then followed by incubation at 37°C at with shaking (200 rpm) for 3 hours. The lysate was then filtered through a 0.22 μm pore size filter and stored at 4 °C until further use. For the *in trans* selection, the lysate was plated on *E. coli* BW25113 *ΔtrxA* strain containing pAG30 or pAG31 vectors that provide wild-type tail fibers.

### 2.9 Confirmation of phage mutants

Following homologous recombination and selection, plaques were picked from plates and resuspended in 1 ml of SM buffer. PCR was used to confirm the presence of phage mutants using primers to differentiate between wildtype and mutant phages (Table S3). PCR products were Sanger sequenced bidirectionally to confirm products were correct. For four phages, whole genome sequencing was carried out. Genomic DNA was extracted from 1 mL of phage lysate using a previously described method [41]. Sequencing libraries were prepared using NexteraXT (Illumina) following the manufacturer’s instructions. Reads were trimmed with sickle using default settings and genomes assembled using SPAdes v3.12.0 [42,43] using ‘-only-assembler’ option and assemblies checked for errors and corrected with pilon [44]. Genomes were annotated with Prokka using phage T7 (accession: V01146) as the source of annotations. SNPs were identified using DNAdiff using default settings [45]. All data was submitted under the project accession PRJEB35760

### 2.10 Type I and Type II CRISPR screening assays

To determine the selection efficiency of the marker based methods (trx and cmk), plaque assays of T7 were performed against *E.coli* BW25113, *E. coli* Δcmk and *E. coli* ΔtrxA with a starting titre of 1 ×10^10^ pfu/ml. The efficiency of plating on *E. coli* Δcmk and *E. coli* ΔtrxA was compared to *E.coli* BW25113. A similar approach was used to determine the efficiency of type I and type II CRISPR systems, with efficiency of plating of phage T7 on strains containing these CRISPR systems compared to *E.coli* BW25113.

## 3. Results

### 3.1. Determination of marker-based (cmk and trxA) selection efficiency against phage T7

In order to test the efficiency of Type I and II CRISPR selection along with the selective markers *trx* and *cmk* all three methods were compared in their ability to select against g17 of wild type T7. To do this the same genetic background was used to try to minimise differences caused by the use of different *E. coli* strains. The strain of choice was *E. coli* BW25113 which allows the Keio collection of mutants to be used[15].

First, the extent to which the absence of host factors reduces T7 progeny was determined. To do this plaque assays were carried out on *E. coli* strains that lacked *cmk* or *trxA*. Efficiency of plating (EOP) on both strains was compared to the control strain *E. coli* BW25113 (Figure 2). In the case of *E. coli* Δ*cmk* cells, the number of progeny phage was reduced by approximately 10^5^-fold, whereas with *E. coli* BW25113 *ΔtrxA* cells no progeny was detected (detection limit ~ 1), reduction by 10^10^-fold (Figure 2).

**Figure 2.**
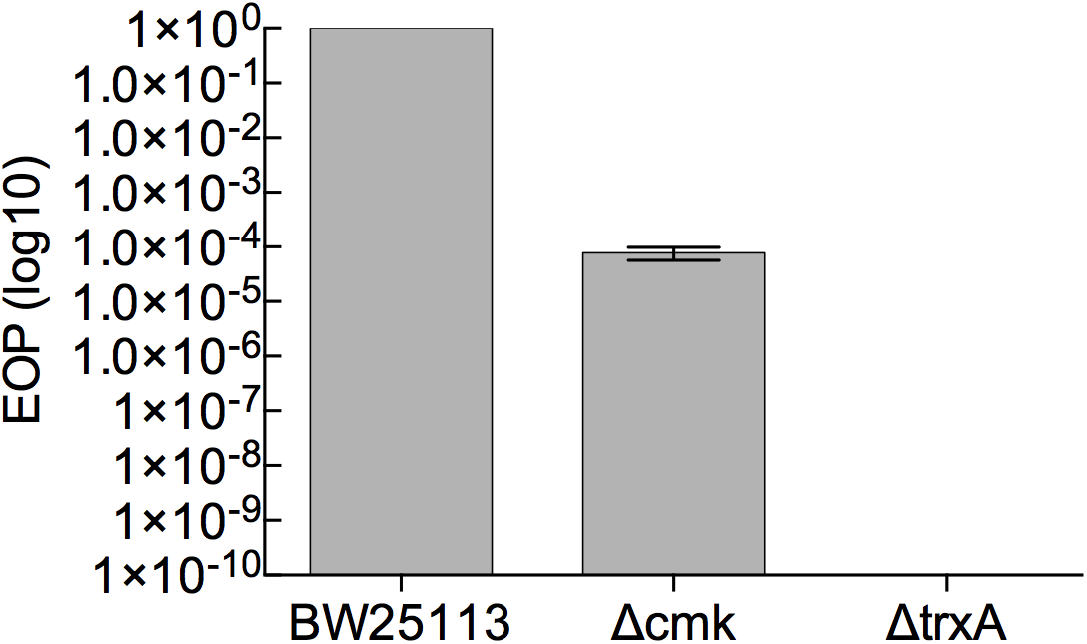
Efficiency of plating for T7 on *E. coli* BW25113, *E. coli* BW25113 Δ*cmk* and *E. coli* BW25113 Δ*trxA* strains. EOP was determined with respect to a reference *E. coli* strain BW25113. EOP data were log10 transformed and are presented as the mean of three independent experiments, n=3. An asterisk indicates that the EOP was below the detection limit which is < 1.

### 3.2 Determination of the efficiency of Type I CRISPR for selecting against wild-type phage T7

A type I CRISPR system that has previously been used to engineer phage T7 was used in this study, and comprises of the vectors pWUR400 and pWUR397 that carry genes encoding for the enzyme cascade and Cas3 respectively [10]. A third plasmid (pAG_1) was constructed in this study that delivers the gRNA and allows for easy swapping of gRNAs to target different regions (Figure 3). It consists of a T7 RNA polymerase promoter at the 5’ of the construct; b) 5’ handle; c) gRNA for a target gene sequence; d) Bbs*I* restriction enzyme sites flanking the gRNA region; e) 3’ handle; f) T7 terminator.

**Figure 3.**
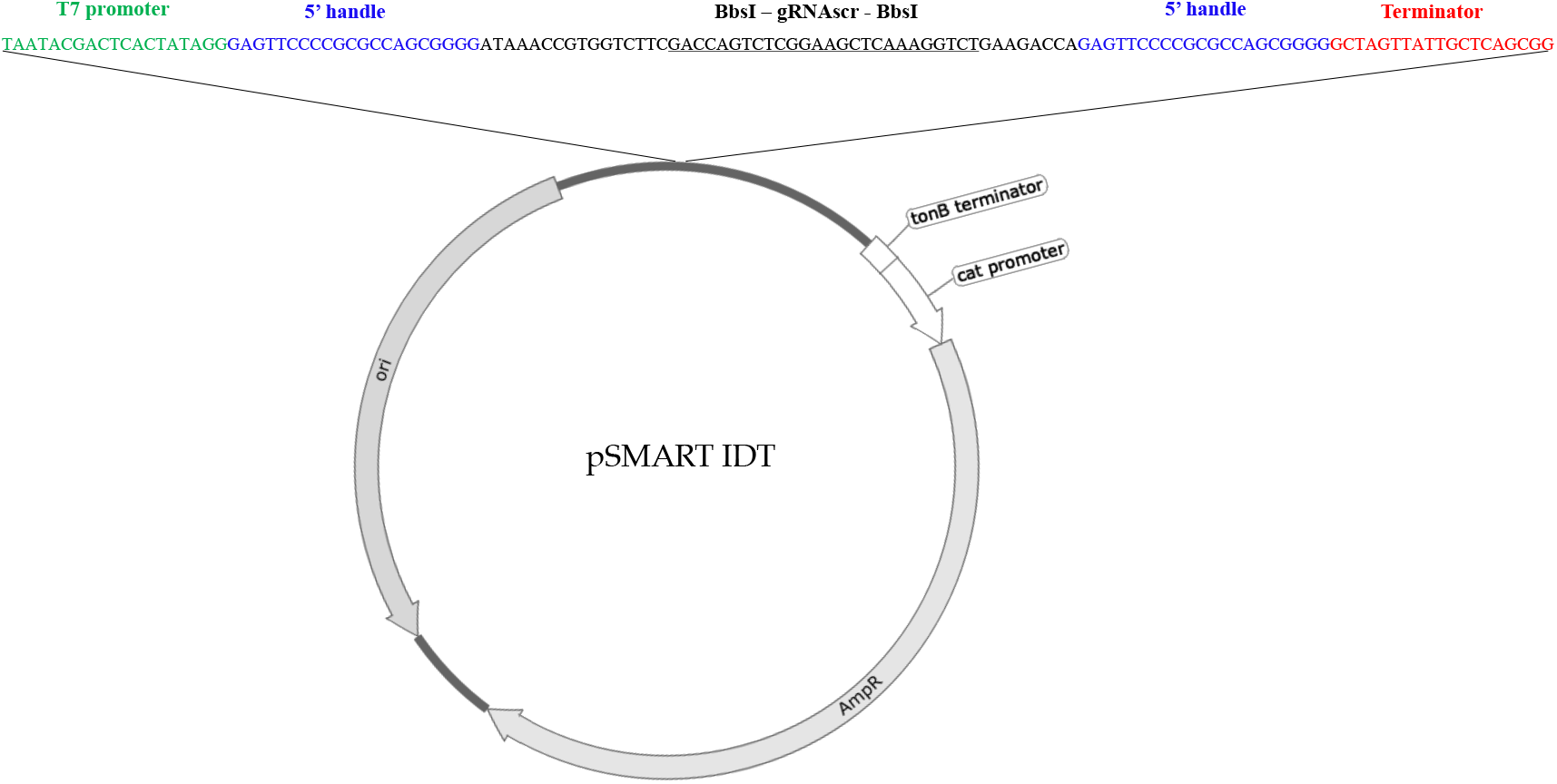
A schematic representation of the vector pAG_1. A 130 bp fragment consisting of a T7 promoter, two handle regions, non-targeting gRNA(scr) with two surrounding BbsI restriction sites and a T7 terminator.

It is known that different gRNAs can alter the efficacy of CRISPR selection [46,47], therefore multiple gRNAS were designed and tested. Eleven different gRNAs, ten targeting *g17* and a scrambled gRNA, were cloned into pAG_1 by Golden Gate cloning (Table S1). The efficiency of each gRNA to target phage DNA was determined by EOP with respect to the control gRNA (**Figure 4.A**). All gRNAs were effective in reducing the number of T7 progeny, with gRNA7 the most effective at reducing T7 progeny by ~90-fold (Figure 4.A). With gRNA2, gRNA5 and gRNA8 reducing T7 progeny by ~10-fold, whereas gRNA1, gRNA6, gRNA9 and gRNA10 showed a reduction of less than 10-fold. The highest reduction in wild type T7 was ~100-fold, which is significantly lower than previous reports of 10,000-fold reduction using this system[25]. No correlation between GC content and EOP was identified between the different gRNAs analysed (Figure 4.B).

**Figure 4.**
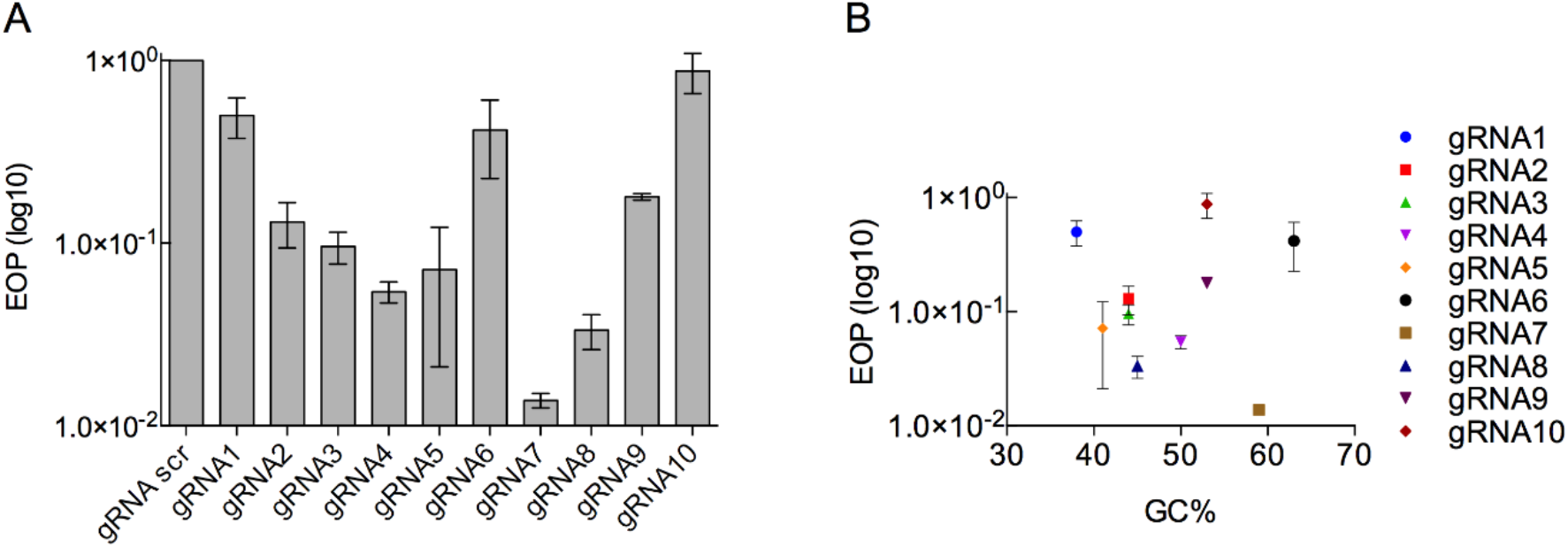
Efficiency of gRNA candidates as marker-less selection system in type I CRISPR-Cas system. (A) Efficiency of plating (EOP) for T7 against *E. coli* BW25113 containing type I CRISPR gRNAs. The (EOP) was determined with respect to a reference *E. coli* strain BW25113/pAG1. EOP data is presented as the mean of three independent experiments. (B) EOP data was plotted against GC% of each gRNA.

*E. coli* BW25113 possesses an innate type I CRISPR system under the control of the transcriptional repressor H-NS (*hns*)[48–50]. The innate CRISPR system is only active if H-NS, is inactivated (Δ*hns*)[49]. The *E. coli* BW25113 used in this study did not contain the *hns* deletion, however, it was necessary to ensure that the innate CRISPR had no effect on the type I CRISPR used here. To verify this *E. coli* BW25113 containing eight gRNAs, including the most efficient gRNA7, was evaluated for its ability to target the T7 genome and accordingly reduce viable T7 progeny. No reduction was identified, ensuring that the introduced CRISPR system used in this study was not overlapping with the innate CRISPR of *E. coli* BW25113 (Figure S1).

A type I CRISPR system has previously been used to target wild-type phage T7, with reductions of two orders of magnitude greater than we obtained [25]. It is possible the different gRNAs are responsible for observed differences in reduction of wild-type T7. However, the previous work of Kiro et al 2014, also induced the expression of the CRISPR system prior to infection with T7. To identify if the pre-induction of a CRISPR system prior to phage addition results in increased efficiency, the same experiment was repeated in strains that contain an inducible T7 RNA polymerase: *E. coli* BL21(DE3) and *E. coli* BL21-AI (S2 and S3). No difference in phage progeny reduction, between induced and un-induced systems, was identified using BL21(DE3) or BL21-AI strains (Figure S3,S4). The carriage of all the plasmids necessary for the type I CRISPR, results in substantial change in growth rate with μ increasing from ~28 min (BL21) to > ~39 mins (BL21 pWUR397 pWUR400 pAG2) (Figure S4).

### 3.3. Determination of type II CRISPR efficiency as a method for engineered phage T7 selection

Eleven gRNAs were designed against *g17*, that spanned the length of the gene and the efficiency of each gRNA was determined. All 11 gRNAs resulted in a reduction in T7 progeny compared to the control. The gRNA2 was the most effective, resulting in a 1000-fold reduction compared to the control (Figure 5). Whereas the majority of gRNAs (gRNA5, gRNA7-gRNA11) showed a reduction in progeny of less than 10-fold, with gRNA3, gRNA4 and gRNA6 resulting in more than 10-fold but less than 100-fold reduction. In working with the type II system, we found that there was no significant change in the growth rate of cells containing Cas9 vector versus no vector containing *E. coli* BW25113 (Figure S9).

**Figure 5.**
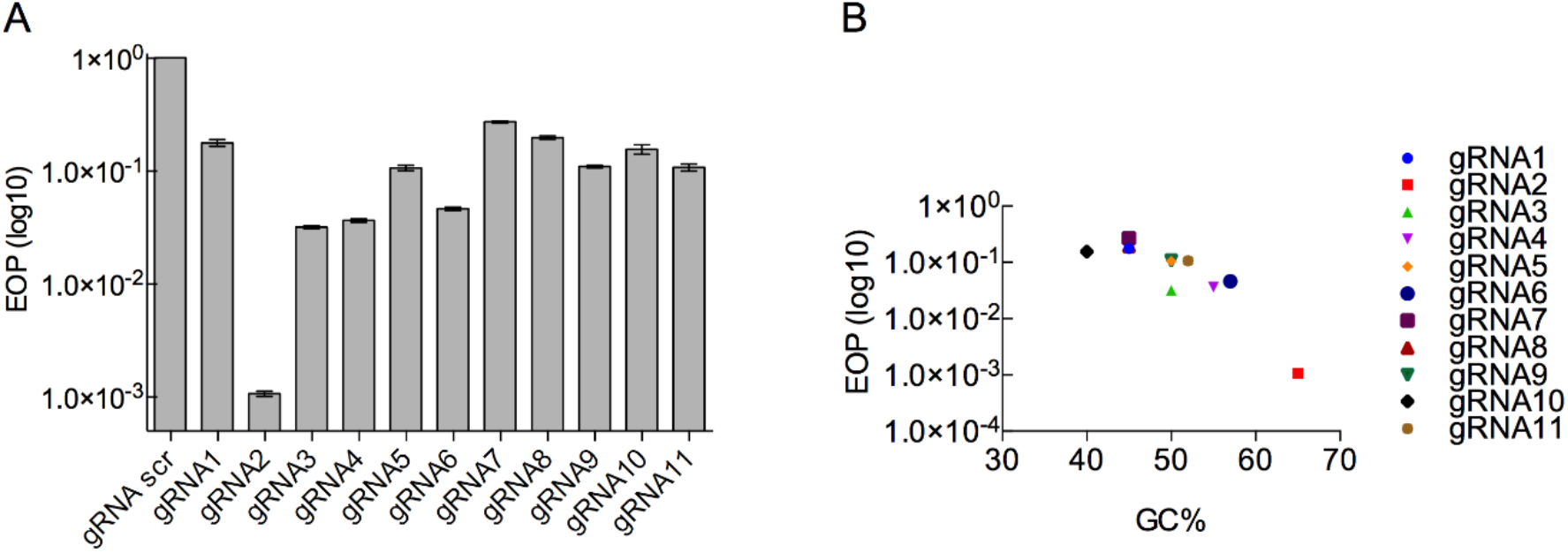
Efficiency of gRNA candidates as marker-less selection system in type II CRISPR-Cas system. (A) Efficiency of plating (EOP) for T7 against *E. coli* BW25113 containing type II CRISPR gRNAs. The EOP was determined with respect to a reference *E. coli* strain BW25113/pAG1. EOP data is presented as the mean of three independent experiments. (B) EOP data plotted against GC% of each gRNA.

### 3.4 Generation of T7 Tail fiber mutants

The use of marker genes was the most efficient at reducing wild type progeny. To check that the method is also effective at selecting phage mutants, we used this method to make a number of mutants that were fusions between the tail fiber of T7 and phages BPP-1 and Yep-phi [51,52]. Phage BPP-1 is very distantly related to phage T7 [53], Yep-phi is more closely related to T7 [52,54]. The tail fiber of BPP-1 contains major tropism determinant (Mtd) domain, where a single mutation allows phage BPP-1 to rapidly evolve and infect the constantly changing receptors of Bordetella pertussis [53]. Although these phages have limited similarity the fusion of the Mtd domain from phage BPP-1 with the T7 tail fiber may allow the host range of phage T7 to be expanded. The tail fiber of Yep-phi (accession: YP_009014859.1) shares 37% protein identity with the T7 homologue, with 53% protein sequence identity over the last 86 amino acids of the C-terminal [54].Thus, it may be more likely to form a functional tail fiber if fused with the T7 version. Eight T7 tail fiber mutants were designed that consisted of domains from both the T7 and BPP-1 tail fiber and one tail fiber mutant that contained domains from T7 and Yep-phi tail fibers (Table S10). This was achieved by first constructing homologous recombination plasmids pAG23 – pAG32(Table S2) with the desired chimeric tail fiber genes and *trxA* for selection. Phage T7 was then used to infect *E. coli* BW25113 Δ*trxA* containing homologous recombination plasmids.

To check that selection with *trxA* was occurring and that chimeric phages could be obtained, phage mutants were created in a two-step manner using an *in trans* tail fiber. Following homologous recombination, the progeny was plated on *E. coli* BW25113 Δ*trxA*, containing a plasmid expressing wild type tail fibers (pAG_30). The resultant phage have a genome which contains a chimeric tail fiber gene, but wild-type tail fibers on the viral particle. Once confirmed as mutants these phages were further propagated on just BW25113 Δ*trxA* to determine if the resultant progeny are functional.

Selection efficiency was measured by PCR screening of plaques to determine the presence of mutants or wild type phage. Between 10 and 30 plaques were screened for each mutant, all of which were positive for phage mutants. Thus, confirming the efficiency of *trxA* marker-based selection for rapidly creating phage mutants. Only one phage mutant was able to infect *E. coli*, the fusion of T7 and Yep-phi tail fibers, with all other phages were non-infective. To ensure the lack of infectivity was due to the non-functional tail fibers and not other deleterious changes in the genome, four phage mutants were sequenced along with the initial wild-type phage T7. In addition to the desired tail fiber mutations, an additional 1-4 SNPs in phages phAG_3, phAG_4 and ph_PM (Table S8). However, these are in regions known to be non-essential for T7 replication and when a wild type tail fiber was provided *in trans* the phage could replicate.

## 4. Discussion

Comparison of selective markers and Type I and II CRISPR systems, revealed that selection with *trxA* was the most efficient method for reducing phage numbers. The use of marker-based selection produced an approximately 10^10^-fold drop in phage progeny, compared to a ~100 x and ~1000x decrease with the most efficient grRNAs using Type I and II CRISPR systems respectively. Previously, the use of *E. coli* Δ*trxA* and *cmk* mutants have been reported to produce an EOP of 10^−2^ and 0.05 respectively [25]. Whereas, within this study, EOPs of 10^−10^ and 10^−4^ was observed. The differences in the observed EOP of plating may simply be due to the growth state of the cells used during a plaque assay. Within this study exponentially growing cells were used, whereas the original work of Kiro *et al* 2014 used cells that had been grown overnight and likely to be in stationary phase of growth[25]. Whilst T7 can infect cells in stationary phase, more productive infections occur in the exponential phase of growth[55]. Thus, when using *E. coli ΔtrxA* or *Δcmk* for positive selection we would suggest infecting *E. coli* in the exponential phase of growth.

The maximum reduction in wild-type T7 progeny, and hence the potential T7 mutant selection efficiency, was found to be 100-fold for the type I CRISPR, when targeting *g17*. This 100-fold reduction was 100 times less than had been achieved by Kiro *et al* when targeting *g1.7* using the same CRISPR system [25]. Although the same type of CRISPR system was used, there were differences in the protocol used as we did not pre-induce the CRISPR system, unlike Kiro *et al* [25]. However, when repeating the experiment using *E. coli* BL21(DE3) and BL21-AI, allowing for induction of the CRISPR system prior to phage addition, we found no difference in Type I CRISPR efficiency with or without induction. Therefore, the difference in observed EOP may simply be down to the selection of the most optimal gRNA. The EOP obtained in this study with different gRNAs varied from 1 to 1 ×10^−4^, which is within the range reported for other studies where the EOP obtained with different gRNAs varies from 1 to 1 ×10^−6^ [29,30].

Using a type II CRISPR system we obtained a 1000-fold reduction when targeting *g17* with gRNA2, which is higher than any gRNA used for the type I system in this study. In targeting *g17* with a similar number of gRNAs distributed across the region of interest, it is clear to see that that type II CRISPR system was generally more efficient than the type I system. The 1000-fold reduction we obtained with a type II CRISPR system is within the range of 10^−1^ and 10^6^ -fold that has previously been reported when using a type II CRISPR system to target phage T4 [29,30]. What gives rise to the large variation in efficiency of different gRNAs is not clear. For non-bacteriophage systems where type II CRISPR has been widely used, there have been numerous studies to predict the efficiency of different gRNAs (see Wilson *et al* 2018 for a review) with large databases of validated gRNAs for different organisms [56,57]. Very high or low GC content of gRNAs has been shown to be an important factor in the decreasing efficiency of gRNAs when targeting human cell lines[58]. Whereas, we found that the efficiency of gRNAs increased with GC content, albeit on small sample. These differences are probably not unexpected given the vastly different backgrounds when using a type II CRISPR system in human cell lines and bacteria.

Given the number of CRISPR systems available to engineer phages, it is often difficult to compare across systems. For instance, the Cas9 used in this study originated from *S. pyogenes* whereas other studies have used Cas9 that originates from *S. thermophiles*, potentially adding to differences in efficiency, combined with the choice of gRNA that is known to be critical [59–61]. Within this study, we directly compared a marker-based system with both type I and II CRISPR systems. Based on the results obtained here, it is clear the most efficient gRNAs for the type II CRISPR system had a higher GC content, an important factor to consider when designing new gRNAs. In contrast with type I system, there was no clear and obvious pattern that may lead to the design of gRNAs that are more effective.

In addition to assessing the effectiveness of a type I and II CRISPR systems and marker-based selection for the ability to target *g17*, we observed differences in ease of use of these systems. In this study we designed a new vector to be used with type I systems that allows gRNAs to be quickly inserted. Whilst this allows the rapid changing of different gRNAs, the system still requires the use of three plasmids. The growth of cells that contain this system was severely inhibited in our hands, making the system difficult and cumbersome to work with compared to a type II system. The choice of method for creating phage mutants will depend on a number of factors and the use of CRISPR allows the creation of clean deletions that cannot be achieved using marker-based methods, whilst also allowing the creation of multiple mutations. Whereas in comparison, marker-based methods are limited to the number of markers that can be selected for. However, the use of marker-based methods has the advantage of being extremely efficient, in particular *trxA*, for the selection of phage mutants and does not require multiple gRNAs to be designed and tested. Within this study we were able to rapidly generate a number of tail fiber mutants (the majority of which ultimately proved to be non-functional in their ability to infect *E. coli)* using *trxA*-based selection. The functional T7-Yep-phi tail fiber hybrid, adds to the growing evidence that functional synthetic phages can be engineered [16,24,25].

## 5. Conclusions

The use of the *trxA* marker-based positive selection proved the most efficient at selecting against wild-type phage T7. We were able to use this system to rapidly engineer eight T7 tail fiber mutants. Although only one of these mutants was ultimately found to be functional. The comparison of type I and II CRISPR systems demonstrated that the type II system was the most efficient at targeting wild type phage T7. This method was also easier to use and would be method of choice if choosing a CRISPR based selection approach. Furthermore, we highlighted that when using CRISPR selection to engineer phage T7 the GC content of the spacer can impact the efficiency. The direct comparison of these three methods provides a basis for further T7 engineering, and aid others who wish to engineer phages in choosing the most appropriate method.

## Author Contributions

The work was conceptualized by A.M.G and A.J. The methodology was carried out by A.M.G, C.H, P.R.M, A.M.B, M.T, J.D, A.P.S. Analysis was carried out by A.M.G, C.H, and A.M. Supervision was carried out by A.M, A.J and C.C. An initial draft was written by A.M.G. All authors reviewed the final manuscript.

## Funding

A. M. G is funded by a doctoral fellowship from the EPSRC & BBSRC Centre for Doctoral Training in Synthetic Biology (grant EP/L016494/1). P. R. M is funded by a doctoral fellowship from the EPSRC Doctoral Training Centre Molecular Organisation and Assembly in Cells (Grant No. EP/F500378/1). C.H is funded by PhD scholarship from the Dept Genetics and Genome, Leicester. A.M is funded by MRC (MR/L015080/1) and NERC (NE/N019881/1).

## Acknowledgments

We thank Luciano Marraffini for plasmid pCas9, Udi Qimron for plasmids pWUR400 and pWUR397 and Mark J van Raaij for his support with tail fiber fusion design.

