## supplementary materials for "Comparison of CRISPR and marker based methods for the engineering of phage T7"

**Table S1. Bacterial strains used in this study.**

| **Strain /Phage** | **Genotype** | Reference |
|---|---|---|
| *E. coli* BW25113 | BW25113 lacI$_{+}$rrnB$_{T14}$ ΔlacZ$_{WJ16}$ hsdR 514 ΔaraBAD$_{AH33}$ ΔrhaBAD$_{LD78}$ rph-1 Δ(araB–D)567 Δ(rhaD–B)568 ΔlacZ4787(::rrnB-3) hsdR514 rph-1 | [15] |
| *E. coli* MG1655 | K-12 F$_{–}$ λ$_{–}$ ilvG$_{–}$ rfb-50 rph-1 | [62] |
| *E. coli* BW25113 ΔtrxA | F-, Δ(araDaraB) 567, ΔlacZ4787(::rrnB-3), λ-, rph-1, ΔtrxA732::kan, Δ(rhaDrhaB)568, hsdR514 | [15] |
| *E. coli* BW25113 Δcmk | F-, Δ(araDaraB)567, ΔlacZ4787(::rrnB-3), λ-, Δcmk-734::kan, rph-1, Δ(rhaDrhaB)568, hsdR514 | [15] |
| *E. coli* BL21-AI | BF$_{–}$ ompT gal dcm lon hsdS$_{B}$(r$_{B–}$m$_{B–}$) [malB$_{+}$]$_{K-12}$(λ$_{S}$) | [63] |
| BL21(DE3) | B F$^{–}$ ompT gal dcm lon hsdS$_{B}$(r$_{B}^{–}$m$_{B}^{–}$) λ(DE3 [lacI lacUV5-T7p07 ind1 sam7 nin5]) [malB$^{+}$]$_{K-12}$(λ$^{S}$) | [64,65] |

**Table S2. Vectors used in this study.**

| Vector | Description Sequence/Relevant Information | Source/ Reference/Notes |
|---|---|---|
| **pWUR400** | Type 1 CRISPR - cas3 under T7 promoter, Kan$^{R}$ | [66] |
| **pWUR397** | Type I cascade genes under T7 promoter, Str$^{R}$ | [66] |
| **pAG_1** | pSMART, Amp$^{R}$, with TAATACGACTCACTATAGGGAGTTCCCCGCGCCAGCGGGGAT AAACCGTGGTCTTCGACCAGTCTCGGAAGCTCAAAGGTCTGA AGACCAGAGTTCCCCGCGCCAGCGGGGCTAGTTATTGCTCA GCGG | This study/used as gRNAscr for type I CRISPR |
| **pAG_2** | pAG1 with TCCTTACGATTAATACAGACTATCGCTTTGCT | This study/ used as gRNA1 for type I CRISPR |
| **pAG_3** | pAG1 with gRNA AAATATTCACGCTAACGGGCGCCTTTACATGA | This study/used as gRNA2 for type I CRISPR |

| | | |
| --- | --- | --- |
| **pAG_4** | pAG1 with gRNA CGGTAACATCCAGTTAGTAGTAAACGGACAGA | This study/used as gRNA3 for type I CRISPR |
| **pAG_5** | pAG1 with gRNA TTACTCGACGTAACTCGATGGTCGTGTAGCCA | This study/ used as gRNA4 for type I CRISPR |
| **pAG_6** | pAG1 with gRNA TACAGTCATTGTTGTTATCTGACCCTCTACCA | This study/used as gRNA5 for type I CRISPR |
| **pAG_7** | pAG1 with gRNA CGTGGACTCAGGTGTGGTCTGGTAGTGCTGGC | This study/used as gRNA6 for type I CRISPR |
| **pAG_8** | pAG1 with gRNA TGTGGTCTGGTAGTGCTGGCGGTGGGGTAAGT | This study/ used as gRNA7 for type I CRISPR |
| **pAG_9** | pAG1 with gRNA ATCTCCGCTTCCGCAATATCTGGATTAAGTGT | This study/ used as gRNA8 for type I CRISPR |
| **pAG_10** | pAG1 with gRNA CTATGAAGTAGATTCCATCGGGGCCAGTACGG | This study/ used as gRNA9 for type I CRISPR |
| **pAG_11** | pAG1 with gRNA TACTGAACGACTGTCTGCAATATTCTTGAATC | This study/ used as gRNA10 for type I CRISPR |
| **pCas_9** | $Cmp^R$, with tgagaccagtctcggaagctcaaaggtctc | [61]/used as gRNAscr for type II CRISPR |
| **pAG_12** | pAG1 with gRNA AAGTGTGACTGTTTCACAGG | This study/ This study/ used as gRNA1 for type II CRISPR |
| **pAG_13** | pAG1 with gRNA AGGCGTGGACTCAGGTGTGG | This study/ used as gRNA2 for type II CRISPR |
| **pAG_14** | pAG1 with gRNA AGTGTGCCAACAACTCTTGG | This study/ used as gRNA3 for type II CRISPR |
| **pAG_15** | pAG1 with gRNA TTCCGCTGCGCATCAATCTG | This study/ used as gRNA4 for type II CRISPR |
| **pAG_16** | pAG1 with gRNA ACGCTACGAACACAAAGCAG | This study/ used as gRNA5 for type II CRISPR |
| **pAG_17** | pAG1 with gRNA CAGCATCCGCTAACTCTGCTC | This study/ used as gRNA6 for type II CRISPR |

| | | |
|---|---|---|
| **pAG_18** | pAG1 with gRNA TACAGTTCCGTAATGAGGCT | This study/ used as gRNA7 for type II CRISPR |
| **pAG_19** | pAG1 with gRNA GGTAAGTGTGACTGTTTCAC | This study/ used as gRNA8 for type II CRISPR |
| **pAG_20** | pAG1 with gRNA GATCTCCGCTTCCGCAATAT | This study/ used as gRNA9 for type II CRISPR |
| **pAG_21** | pAG1 with gRNA CTTCCGCAATATCTGGATTA | This study/ used as gRNA10 for type II CRISPR |
| **pAG_22** | pAG1 with gRNA AATACACTCCAACGGTCTCG | This study/ used as gRNA11 for type II CRISPR |
| **pSB6A1** | $Amp^R$, | Registry of Standard Biological parts |
| **pAG_23** | pSB6A1 with HR1*, 513-1146 (*mtd), RBS (B0034), trxA (full sequence),* HR2** | This study |
| **pAG_24** | pSB6A1 with HR1*, 513-1146 (*mtd), RBS (B0034), trxA (full sequence),* HR2** | This study |
| **pAG_25** | pSB6A1 with HR1*, 513-1146 (*mtd), RBS (B0034), trxA (full sequence),* HR2** | This study |
| **pAG_26** | pSB6A1 with HR1*, 513-1146 (*mtd), RBS (B0034), trxA (full sequence),* HR2** | This study |
| **pAG_27** | pSB6A1 with HR1*, 513-1146 (*mtd), RBS (B0034), trxA (full sequence),* HR2** | This study |
| **pAG_28** | pSB6A1 with HR1*, 513-1146 (*mtd), RBS (B0034), trxA (full sequence),* HR2** | This study |
| **pAG_29** | pSB6A1 with HR1*, 513-1146 (*mtd), RBS (B0034), trxA (full sequence),* HR2** | This study |
| **pAG_30** | pSB6A1 with full *g17* under T7 promoter | This study |
| **pAG_31** | pSEVA551 with codon optimised full *g17* under T7 promoter | This study |
| **pAG_32** | pSB6A1 with HR1*, 1455-1707 (Yep-phi, *g17*), *RBS (B0034), trxA (full sequence),* HR2** | This study |

HR1* gccgttgtggccactgatggtaatattcaaggtactaagtggggaggtaaatggctggatgcttacctacgtgacagcttcgttgcgaag

HR2** ttggtaaatcacaaggaaagacgtgtagtccacggatggactctcaaggaggtacaaggtgctatcattagactttaacaacgaattgat

**Table S3. Primers used in this study.**

| Primer Sequence 5' to 3' | Use |
|---|---|
| ACCGTCCTTACGATTAATACAGACTATCGCTTTGCT | Forward primer to generate insert for pAG_2 |

| | |
|---|---|
| ACTCAGCAAAGCGATAGTCTGTATTAATCGTAAGGA | Reverse primer to generate insert for pAG_2 |
| ACCGAAATATTCACGCTAACGGGCGCCTTTACATGA | Forward primer to generate insert for pAG_3 |
| ACTCTCATGTAAAGGCGCCCGTTAGCGTGAATATTT | Reverse primer to generate insert for pAG_3 |
| ACCGCGGTAACATCCAGTTAGTAGTAAACGGACAGA | Forward primer to generate insert for pAG_4 |
| ACTCTCTGTCCGTTTACTACTAACTGGATGTTACCG | Reverse primer to generate insert for pAG_4 |
| ACCGTTACTCGACGTAACTCGATGGTCGTGTAGCCA | Forward primer to generate insert for pAG_5 |
| ACTCTGGCTACACGACCATCGAGTTACGTCGAGTAA | Reverse primer to generate insert for pAG_5 |
| ACCGTACAGTCATTGTTGTTATCTGACCCTCTACCA | Forward primer to generate insert for pAG_6 |
| ACTCTGGTAGAGGGTCAGATAACAACAATGACTGTA | Reverse primer to generate insert for pAG_6 |
| ACCGCGTGGACTCAGGTGTGGTCTGGTAGTGCTGGC | Forward primer to generate insert for pAG_7 |
| ACTCGCCAGCACTACCAGACCACACCTGAGTCCACG | Reverse primer to generate insert for pAG_7 |
| ACCGTGTGGTCTGGTAGTGCTGGCGGTGGGGTAAGT | Forward primer to generate insert for pAG_8 |
| ACTCACTTACCCCACCGCCAGCACTACCAGACCACA | Reverse primer to generate insert for pAG_8 |
| ACCGATCTCCGCTTCCGCAATATCTGGATTAAGTGT | Forward primer to generate insert for pAG_9 |
| ACTCACACTTAATCCAGATATTGCGGAAGCGGAGAT | Reverse primer to generate insert for pAG_9 |
| ACCGCTATGAAGTAGATTCCATCGGGGCCAGTACGG | Forward primer to generate insert for pAG_10 |
| ACTCCCGTACTGGCCCCGATGGAATCTACTTCATAG | Reverse primer to generate insert for pAG_10 |
| ACCGTACTGAACGACTGTCTGCAATATTCTTGAATC | Forward primer to generate insert for pAG_11 |
| ACTCGATTCAAGAATATTGCAGACAGTCGTTCAGTA | Reverse primer to generate insert for pAG_11 |
| AAACAAGTGTGACTGTTTCACAGG | Forward primer to generate insert for pAG_12 |
| AAAACCTGTGAAACAGTCACACTT | Reverse primer to generate insert for pAG_12 |
| AAACAGGCGTGGACTCAGGTGTGG | Forward primer to generate insert for pAG_13 |
| AAAACCACACCTGAGTCCACGCCT | Reverse primer to generate insert for pAG_13 |
| AAACAGTGTGCCAACAACTCTTGG | Forward primer to generate insert for pAG_14 |

| | |
|---|---|
| AAAACCAAGAGTTGTTGGCACACT | Reverse primer to generate insert for pAG_14 |
| AAACGTTCCGCTGCGCATCAATCTG | Forward primer to generate insert for pAG_15 |
| AAAACAGATTGATGCGCAGCGGAAC | Reverse primer to generate insert for pAG_15 |
| AAACGACGCTACGAACACAAAGCAG | Forward primer to generate insert for pAG_16 |
| AAAACTGCTTTGTGTTCGTAGCGTC | Reverse primer to generate insert for pAG_16 |
| AAACGAGCAGAGTTAGCGGATGCTG | Forward primer to generate insert for pAG_17 |
| AAAACAGCATCCGCTAACTCTGCTC | Reverse primer to generate insert for pAG_17 |
| AAACGAGCCTCATTACGGAACTGTA | Forward primer to generate insert for pAG_18 |
| AAAATACAGTTCCGTAATGAGGCTC | Reverse primer to generate insert for pAG_18 |
| AAACGGTAAGTGTGACTGTTTCAC | Forward primer to generate insert for pAG_19 |
| AAAAGTGAAACAGTCACACTTACC | Reverse primer to generate insert for pAG_19 |
| AAACGATCTCCGCTTCCGCAATATC | Forward primer to generate insert for pAG_20 |
| AAAAGATATTGCGGAAGCGGAGATC | Reverse primer to generate insert for pAG_20 |
| AAACGTAATCCAGATATTGCGGAAG | Forward primer to generate insert for pAG_21 |
| AAAACTTCCGCAATATCTGGATTAC | Reverse primer to generate insert for pAG_21 |
| AAACGCGAGACCGTTGGAGTGTATT | Forward primer to generate insert for pAG_22 |
| AAAAAATACACTCCAACGGTCTCGC | Reverse primer to generate insert for pAG_22 |
| GTGACAGCTTCGTTGCGAAGATCAAGTTTCGCCCGGCTGC | For pAG_26 construct Gibson assembly; *mtd* g-block forward |
| TGACCTCCTTAAAGTAAATCACAAAAAACCCCTAGCCGCC | *mtd* g-block reverse |
| ACGCCAACCTGGCTTAATGATTGGTAAATCACAAGGAAAGACG | pSB6A1 (Yep-phi) *p17*g-block g-block forward |
| GCAGCCGGGCGAAACTTGATCTTCGCAACGAAGCTGTCAC | pSB6A1 (Yep-phi) *p17*g-block g-block reverse |
| GGCGGCTAGGGGTTTTTTGTGATTTACTTTAAGGAGGTCAAATGAG | *trxA* g-block forward |
| CTTTCCTTGTGATTTACCAATCATTAAGCCAGGTTGGCGT | *trxA* g block reverse |

| | |
|---|---|
| GTGACAGCTTCGTTGCGAAGAACGAATACAGCCTGTGGGA | For pAG_25 construct Gibson assembly; *mtd* g-block forward |
| TGACCTCCTTAAAGTAAATCACAAAAAACCCCTAGCCGCC | *mtd* g-block reverse |
| ACGCCAACCTGGCTTAATGATTGGTAAATCACAAGGAAAGACG | pSB6A1 (Yep-phi) *p17*g-block g-block forward |
| TCCCACAGGCTGTATTCGTTCTTCGCAACGAAGCTGTCAC | pSB6A1 (Yep-phi) *p17*g-block g-block reverse |
| GGCGGCTAGGGGTTTTTTGTGATTTACTTTAAGGAGGTCAAATGAG | *trxA* g-block forward |
| CTTTCCTTGTGATTTACCAATCATTAAGCCAGGTTGGCGT | *trxA* g-block reverse |
| GTGACAGCTTCGTTGCGAAGAACGAATACAGCCTGTGGGA | For pAG_24 construct Gibson assembly; *mtd* g-block forward |
| TGACCTCCTTAAAGTAAATCACAAAAAACCCCTAGCCGCC | to amplify *mtd* g-block reverse |
| ACGCCAACCTGGCTTAATGATTGGTAAATCACAAGGAAAGACG | pSB6A1 (Yep-phi) *p17*g-block g-block forward |
| TCGGCCTTGATGAAAGCCGTCTTCGCAACGAAGCTGTCAC | pSB6A1 (Yep-phi) *p17*g-block g-block reverse |
| GGCGGCTAGGGGTTTTTTGTGATTTACTTTAAGGAGGTCAAATGAG | *trxA* g-block forward |
| CTTTCCTTGTGATTTACCAATCATTAAGCCAGGTTGGCGT | *trxA* g-block reverse |
| ACCACCTGATTCTTGAGTAGCGGGGCCGAAAGGCCCCGCC | For pAG_27 construct Gibson assembly; pAG_24 forward |
| ACCAGATCGTGCCGCGCCAGCTTCGCAACGAAGCTGTCACGTAGGTAAGC | pAG24 reverse |
| GTGACAGCTTCGTTGCGAAGCTGGCGCGGCACGATCTGGT | *mtd* g-block forward |
| GGCGGGGCCTTTCGGCCCCGCTACTCAAGAATCAGGTGGTCACAGACG | *mtd* g-block reverse |
| GTGACAGCTTCGTTGCGAAGAAGTTTCGCCCGGCTGCGCT | For pAG_23 construct assembly; *mtd* g-block forward |
| TGACCTCCTTAAAGTAAATCCGGTTGCCTTGGCGGGGCCT | *mtd* g-block reverse |
| ACGCCAACCTGGCTTAATGATTGGTAAATCACAAGGAAAGACG | pSB6A1 (Yep-phi) *p17*g-block g-block forward |
| AGCGCAGCCGGGCGAAACTTCTTCGCAACGAAGCTGTCAC | pSB6A1 (Yep-phi) *p17*g-block g-block reverse |

| | |
|---|---|
| AGGCCCCGCCAAGGCAACCGGATTTACTTTAAGGAGGTCAAATGAG | *trxA* g-block forward |
| CTTTCCTTGTGATTTACCAATCATTAAGCCAGGTTGGCGT | *trxA* g-block reverse |
| ACCACCTGATTCTTGAGTAGCGGGGCCGAAAGGCCCCGCC | For pAG_28 construct Gibson assembly; pAG_24 forward |
| ATGAAAGCCGTTTTCACCAGCTTCGCAACGAAGCTGTCACGTAGGTAAGC | pAG_24 reverse |
| GTGACAGCTTCGTTGCGAAGCTGGTGAAAACGGCTTTCAT | *mtd* g-block forward |
| GGCGGGGCCTTTCGGCCCCGCTACTCAAGAATCAGGTGGT | *mtd* g-block reverse |
| GGCCCTTTCGTCTTCAAGAATGTTAACTTGAGGGAGCGTA | *trxA* g-block forward |
| CACGATGCGTCCGGCGTAGATCAAATCAATTCGTTGTTAAAGTC | *trxA* g-block reverse |
| TTAACAACGAATTGATTTGATCTACGCCGGACGCATCGTG | pSB6A1 forward |
| ACGCTCCCTCAAGTTAACATTCTTGAAGACGAAAGGGCCTC | pSB6A1 reverse |
| GGCCTGCAGGAGTCACTACTAGTAGCGGCCGCTG | Gibson primers to generate pAG_32 |
| CGCCGCGCGCCCCGAAGTTAGTTTCGAACTAAGATTTGC | |
| TCTTAGTTCGAAACTAACTTCGGGGCGCGCGGCG | |
| CAGCGGCCGCTACTAGTAGTGACTCCTGCAGGCCTTAATCAATT CGTTGTT | |
| TGGACTCACAAAGAAAAAACGCCCGGTGTGCAAGACCGAGCGTTCTGAACAATT ACTCGTTCTCCACCATGATTGC | Gibson primers to generate pAG_30, *g17* forward |
| CTCACTATAGGGAGAACTAGAGAAAGAGGAGAAATACTAGATGGCTAACGTAAT TAAAACCGTTTTGAC | *g17* reverse |
| GTCAAAACGGTTTTAATTACGTTAGCCATCTAGTATTTCTCCTCTTTCTCTAGTTCT CCCTATAGTGAG | pSB6A1 forward |
| GCAATCATGGTGGAGAACGAGTAATTGTTCAGAACGCTCGGTCTTGCACACCGG GCGTTTTTTCTTTGTGAGTCCA | pSB6A1 reverse |
| TCGGCTGGCTTTGTGGCTAACG | T7 sequencing, before *g17* |
| ACCTCCTTGAGAGTCCATCCGTGG | T7 sequencing, after *g17* |

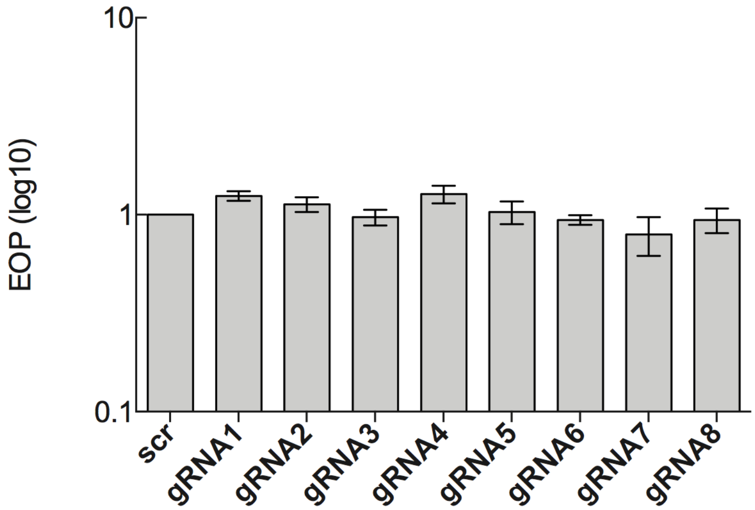

**Figure S1. Efficiency of gRNAs only vectors of type I CRISPR-Cas system**. Efficiency of plating for T7 against *E. coli* BW25113 containing type I CRISPR gRNAs only. The T7 efficiency of plating was determined with respect to a reference *E. coli* BW25113/pAG1 strain. EOP data is presented as the mean of three independent experiments. The concentration of the phage stock added $2x10^9$ PFU/ml.

*E. coli* BL21-AI strain containing Type I CRISPR-Cas plasmids were grown in liquid LB medium to $OD_{600}$ = 0.3-0.4. Expression of the CRISPR-Cas genes was induced by the addition of IPTG to a final concentration of 0.1 mM and the cultures incubated at 37$^o$C for 1 hour. Each of the induced cultures were then used to carryout plaque assays, followed by incubation at 37$^o$C overnight to calculate efficiency of plating.

*E. coli* BL21(DE3) strain containing the Type I CRISPR-Cas plasmids were grown in liquid LB medium to $OD_{600}$ = 0.3-0.4. Expression of the CRISPR-Cas genes was induced by the addition of IPTG to a final concentration of 0.1 mM and the cultures incubated at 37$^o$C for 1 hour. For each of the induced cultures, 1 mL was combined with 8 mL of 0.7% LB agar, with and without 0.1 mM IPTG, and plated onto 1.5% LB agar plates before being further incubated at 37$^o$C for 1 hour. T7 was used to perform spot assays followed by incubation at 37$^o$C overnight to calculate efficiency of plating.

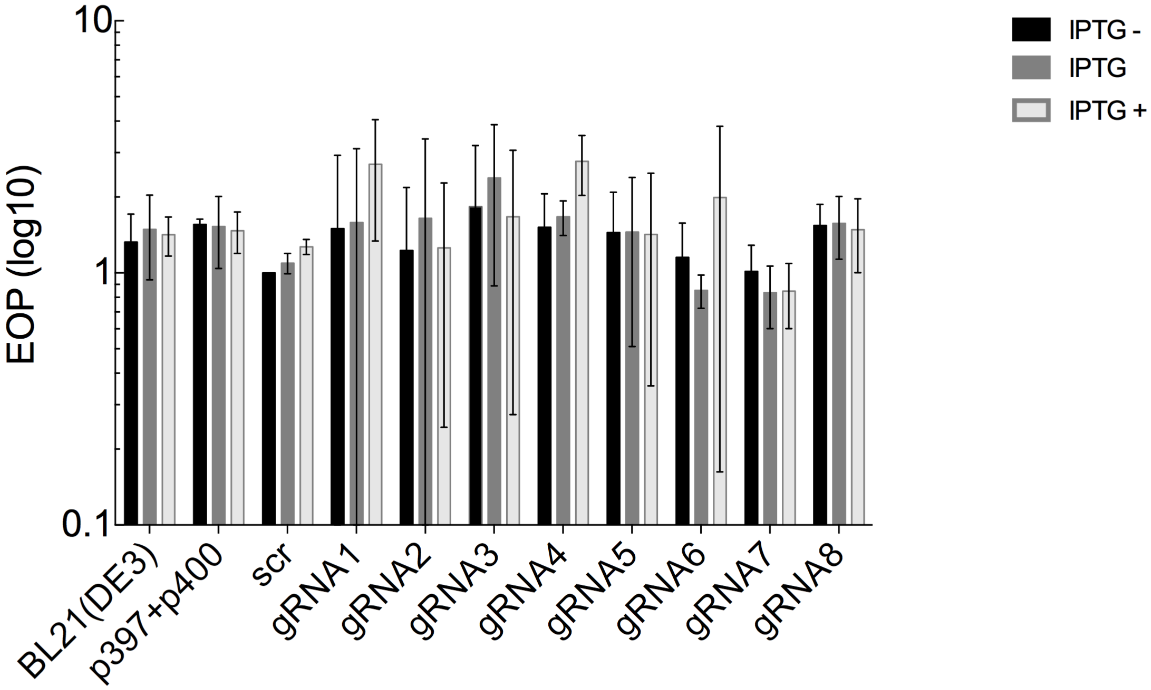

**Figure S2. Efficiency of type I CRISPR-Cas system with and without IPTG induction.** Efficiency of plating for T7 against *E. coli* BL21 (DE3) containing pWUR400 and pWUR397, and one of the nine gRNAs. The T7 efficiency of plating was determined with respect to a reference uninduced *E. coli* BL21(DE3) containing gRNA(scr). IPTG (-) group has no IPTG added. IPTG group had IPTG added only to the bacterial cultures. IPTG (+) group had IPTG added to the bacterial cultures as well as the top agar. EOP data is presented as the mean of three independent experiments. The concentration of the phage stock added $10^{10}$ PFU/ml.

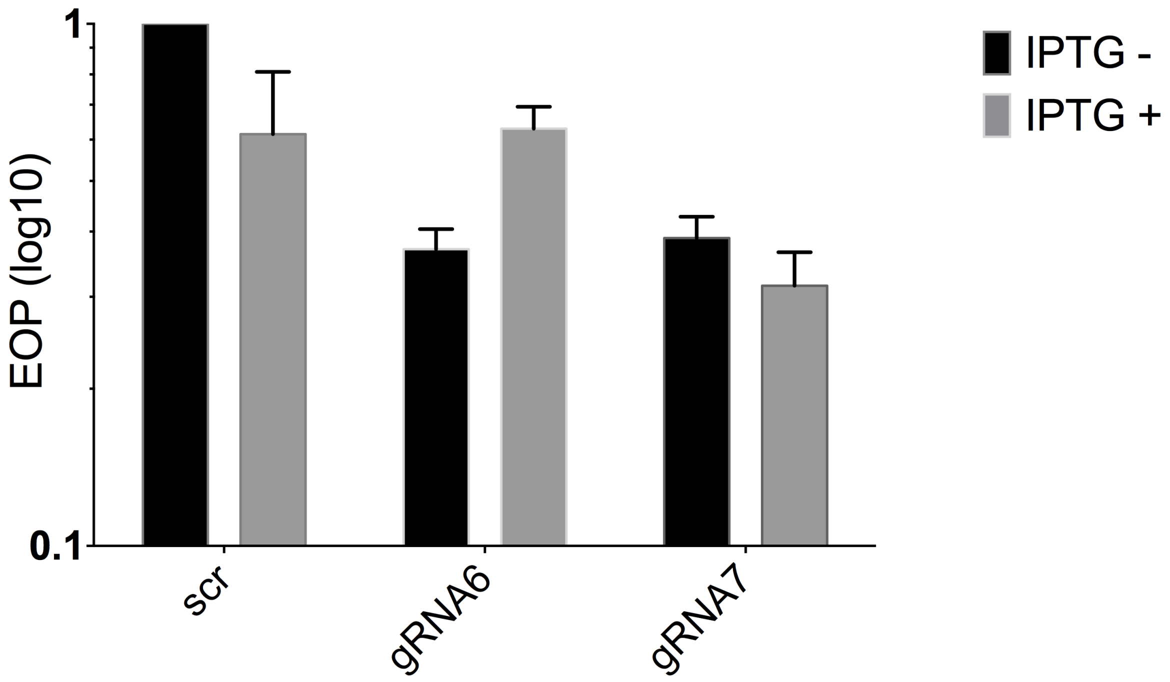

**Figure S3. Efficiency of type I CRISPR-Cas system with and without IPTG induction.** Efficiency of plating for T7 against *E. coli* BW25113 containing pWUR400 and pWUR397 and one of the three gRNA vectors pAG1 (gRNAscr), pAG7 (gRNA6) and pAG8 (gRNA7). The T7 efficiency of plating was determined with respect to a reference undinduced *E. coli* BW25113/pAG1 strain. EOP data is presented as the mean of three independent experiments. The concentration of the phage stock added 2x10$^{9}$PFU/ml.

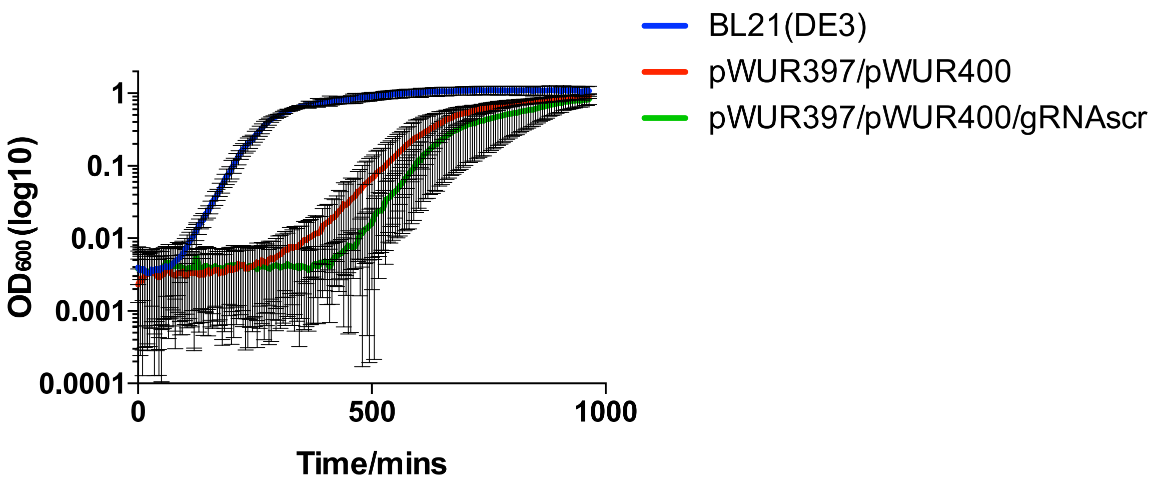

**Figure S4. Growth of BL21(DE3) with and without type I CRISPR vectors**. The data is presented for twenty-four technical replicates. The growth rate for each of the strains was determined. The growth rate for BL21(DE3) was 27.97 (+/- 0.46) mins. The growth rate for BL21(DE3) containing pWUR397/pWUR400 vectors was 38.92 (+/- 0.80) mins. The growth rate for BL21(DE3) containing pWUR397/pWUR400/gRNAscr was 96.14 (+/- 92.68) mins.

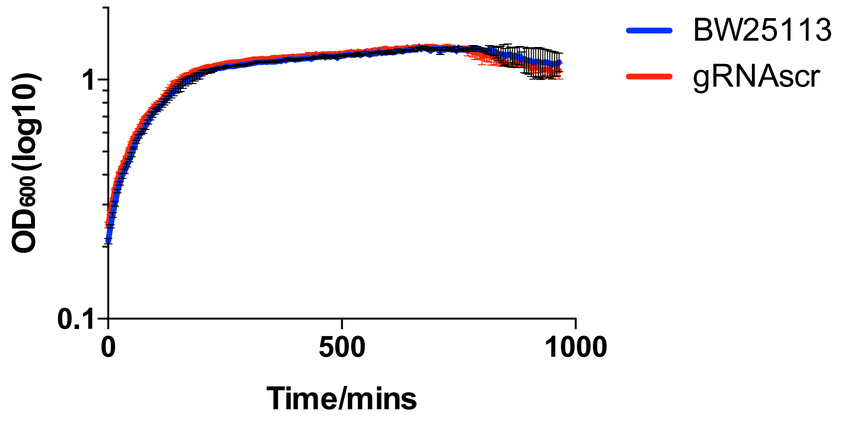

**Figure S5. Growth of BW25113 with and without type II CRISPR vector.** The data is presented for three technical replicates.

**Table S4. Summary of phage mutants generated in this study.**

| T7 and BPP-1 tail fibre fusions | Gp17/*g17* aa/bp | Mtd/*mtd* aa/bp | Phage mutants generated* | HR vectors used for phage mutants |
|---|---|---|---|---|
| 1 | 1 - 466/1-1398 | 171 – 382/513 – 1146 | phAG_1 | pAG23 |
| 2 | 1 - 466/1-1398 | 55 – 382/165 – 1146 | phAG_2 | pAG24 |
| 3 | 1 - 466/1-1398 | 163 – 382/489 – 1146 | phAG_3 | pAG25 |
| 4 | 1 - 466/1-1398 | 170 – 382/510 – 1146 | phAG_4 | pAG26 |
| 5 | 1 - 466/1-1398 | 47 – 382/141 – 1146 | phAG_5 | pAG27 |
| 6 | 1 - 466/1-1398 | 52 – 382/156 – 1146 | phAG_6 | pAG28 |
| 7 | N/A | N/A | phAG_7** | pAG29 |
| 8 | 1 - 466/1-1398 | (Yep-phi/gp17) 485-569/1455-1707 | phPM | pAG32 |

*each mutant contains full sequence *trxA* after *mtd*/Yep-phi *g17* sequence insert.

**phAG_7 has g17 replaced with full sequence of *trxA*.

**Table S5. Efficiency of generating phage T7 mutants.** *In trans* method tail fiber mutant efficiency represented as plaque PCR screening output.

| Phage Mutant | Plaques screened | Successful mutants (%) |
|---|---|---|
| **phAG_1** | **30** | **100** |
| **phAG_2** | **30** | **100** |
| **phAG_3** | **30** | **100** |
| **phAG_4** | **15** | **100** |
| **phAG_5** | **15** | **100** |
| **phAG_6** | **10** | **100** |
| **phAG_7** | **10** | **100** |

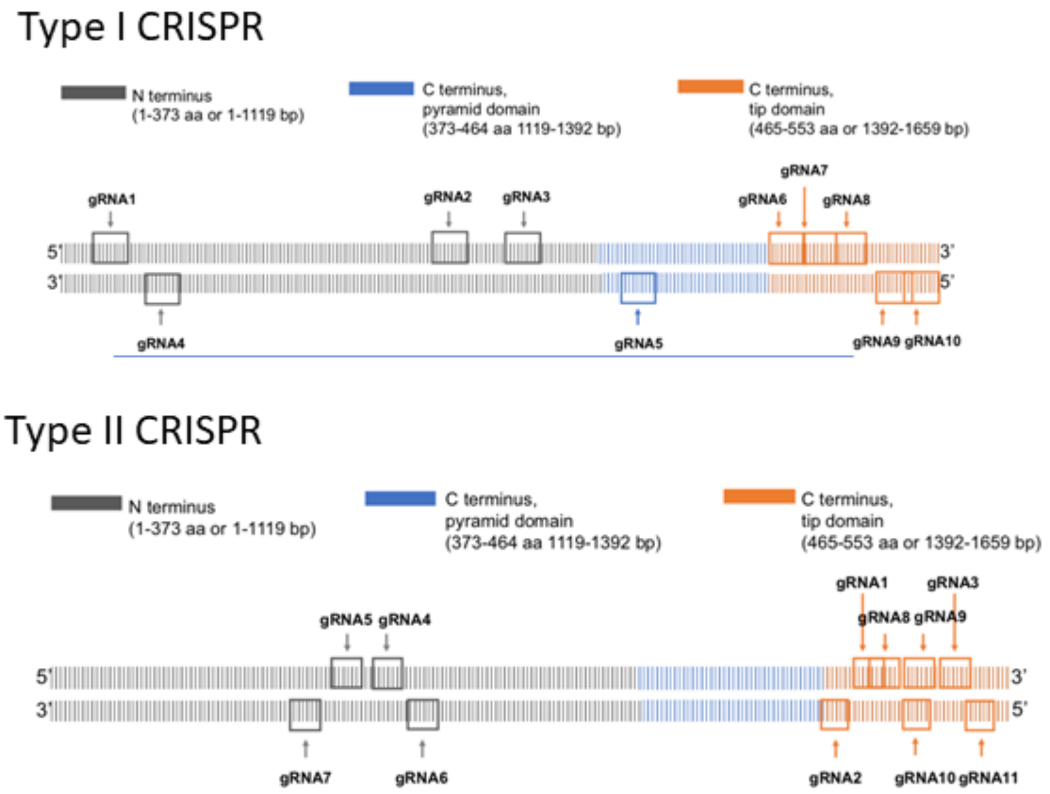

**Figure S6.** Representation of the distribution of gRNAs on *g17* designed for type I and type II CRISPRs.

**Table S6. Single nucleotide variations detected in phage T7 mutants**

| Phage | SNV Position | Wild type | Mutant | Amino Acid change |
|---|---|---|---|---|
| phAG_3 | 24876 | A | G | K->R |
| phAG_4 | 742 | T | C | intergenic |
| phAG_4 | 1897 | C | . | D->E |
| phAG_4 | 1899 | A | C | Q->K |
| phAG_4 | 1900 | G | A | Q->K |
| phAG_4 | 19560 | T | G | V->G |
| phPM | 26041 | A | G | I>M |

**Figure S7. Sequences of g-blocks used in this study.**

(BPP-1) *mtd* g-block:

ATGAGTACCGCAGTCCAATTCCGAGGTGGAACGACCGCCCAGCACGCAACGTTCACGGGCGCCGCCCGTGA
GATTACCGTCGATACCGACAAGAACACGGTCGTTGTGCATGACGGTGCTACCGCTGGCGGCTTCCCCCTGGC
GCGGCACGATCTGGTGAAAACGGCTTTCATCAAGGCCGACAAGTCGGCCGTCGCCTTCACGCGCACCGGCA
ACGCAACGGCCAGCATCAAGGCTGGCACCATCGTGGAGGTCAACGGCAAGCTGGTGCAGTTCACCGCCGAC
ACGGCCATCACCATGCCGGCGCTGACGGCCGGCACCGACTACGCCATCTACGTCTGCGACGATGGCACGGT
GCGCGCCGATTCCAACTTTTCGGCGCCCACTGGCTACACCTCGACCACGGCGCGCAAGGTGGGCGGCTTCCA
CTATGCGCCGGGAAGCAACGCTGCAGCGCAGGCTGGTGGAAACACCACGGCGCAGATCAACGAATACAGC
CTGTGGGACATCAAGTTTCGCCCGGCTGCGCTCGACCCGCGCGGCATGACGCTGGTTGCCGGCGCGTTTTGG
GCAGACATCTATCTGCTAGGCGTCAACCACCTGACCGATGGCACCAGCAAATACAACGTGACAATTGCAGA
TGGTAGTGCATCACCTAAGAAATCTACCAAGTTCGGTGGAGACGGCAGCGCGGCCTACAGTGACGGAGCTT
GGTACAACTTCGCTGAGGTCATGACTCATCACGGTAAGCGCCTGCCTAACTACAACGAATTCCAGGCGCTGG
CTTTCGGCACGACCGAGGCTACGTCCAGCGGCGGCACCGACGTGCCCACCACCGGCGTGAACGGCACGGGC
GCAACGAGCGCGTGGAACATCTTCACGTCCAAGTGGGGCGTTGTGCAGGCGTCCGGTTGCTTGTGGACGTGG
GGTAACGAGTTCGGCGGCGTGAATGGCGCATCCGAATACACGGCCAACACTGGCGGCAGAGGATCGGTGTA
CGCCCAGCCCGCTGCTGCGCTATTCGGCGGCGCCTGGAACGGCACGTCGCTCTCGGGTTCTCGCGCTGCGCT
CTGGTACAGCGGGCCGTCGTTCTCGTTCGCGTTCTTCGGGGCGCGCGGCGTCTGTGACCACCTGATTCTTGAG
TAGCGGGGCCGAAAGGCCCCGCCAAGGCAACCGTACTCGAACCCCTAGCCCGCTCTTATCGGGCGGCTAGG
GGTTTTTTGT

HR1-*trxA*-HR2 g-block:

TGTTAACTTGAGGGAGCGTAGGAAATAATACGACTCACTATAGGGAGAGGCGAAATAATCTTCTCCCTGTAG
TCTCTTAGATTTACTTTAAGGAGGTCAAATGAGCGATAAAATCATTCACCTGACCGATGACTCTTTTGATACC
GACGTGCTGAAAGCTGATGGTGCAATTCTGGTTGATTTCTGGGCAGAGTGGTGCGGCCCTTGCAAAATGATC
GCTCCAATCCTGGACGAAATTGCGGACGAATATCAGGGTAAGCTGACTGTGGCCAAACTGAACATTGACCA
GAACCCTGGCACCGCACCGAAATACGGTATCCGTGGCATCCCAACTCTGCTGCTGTTCAAAAACGGTGAAGT
GGCAGCAACCAAAGTAGGCGCTCTGTCTAAAGGCCAACTGAAAGAGTTCCTGGACGCCAACCTGGCTTAAT

GATTGGTAAATCACAAGGAAAGACGTGTAGTCCACGGATGGACTCTCAAGGAGGTACAAGGTGCTATCATT
AGACTTTAACAACGAATTGATTTGA

(Yep-phi) *g17* g-block:

GCCGTTGTGGCCACTGATGGTAATATTCAAGGTACTAAGTGGGGAGGTAAATGGCTGGATGCTTACCTACGT
GACAGCTTCGTTGCGAAGAGCTCTGGTTGGACTGAGGTATGGCAAGGCTCTGCTGGTGGTGGTGTTTCAGTA
AGCCTCTCACAGGATGTCCGCTGGAGAACTATCTGGATTCTAGCTAATAATGGCATGTGTTCTGTTCAGATTG
GAGCTGATGCTACTTACTTCATGGTGGTTATGGGTGGTTGGTTGAAGTTCACAATTTCCAACAACGGGAGAA
CTTTCCGTAACGACCAAGATCGAAATACAGTACCTGAGCAAATCTTAGTTCGAAACTAATAATTGGTAAATC
ACAAGGAAAGACGTGTAGTCCACGGATGGACTCTCAAGGAGGTACAAGGTGCTATCATTAGACTTTAACAA
CGAATTGATA
